# AlmostSignificant: Simplifying quality control of high-throughput sequencing data

**DOI:** 10.1101/053702

**Authors:** Joseph Ward, Christian Cole, Melanie Febrer, Geoffrey Barton

## Abstract

**Motivation:** The current generation of DNA sequencing technologies produce a large amount of data quickly. All of these data need to pass some form of quality control processing and checking before they can be used for any analysis. The large number of samples that are run through Illumina sequencing machines makes the process of quality control an onerous and time-consuming task that requires multiple pieces of information from several sources.

**Results:** AlmostSignificant is an open-source platform for aggregating multiple sources of quality metrics as well as meta-data associated with DNA sequencing runs from Illumina sequencing machines. AlmostSignificant is a graphical platform to streamline the quality control of DNA sequencing data, to collect and store these data for future reference and to collect extra meta-data associated with the sequencing runs to check for errors and monitor the volume of data produced by the associated machines. AlmostSignificant has been used to track the quality of over 80 sequencing runs covering over 2500 samples produced over the last three years.

**Availability:** The code and documentation for AlmostSignificant is freely available at https://github.com/bartongroup/AlmostSignificant.

## 1 Introduction

The current generation of high-throughput, short-read DNA sequencing technologies quickly generate large datasets, but all sequencing technologies produce errors, whether systematic, technical, or simple human mistakes. Tracking the quality of the sequence data is critical to a successful experiment, but can become time consuming for all but the smallest projects. Monitoring the success of a sequencing run requires dedicated software located on the sequencing machine or standalone software provided by its manufacturer. These are often used in conjunction with tools such as FastQC (www.bioinformatics.babraham.ac.uk/projects/fastqc) for assessing the quality of the output fastq files and FastQScreen (www.bioinformatics.babraham.ac.uk/projects/fastq_screen) to check for contaminating DNA. Although these tools are simple to use and interpret for a handful of samples or a single project, this is difficult for sequencing facilities that handle many sequencing runs simultaneously.

AlmostSignificant is open-source software designed to simplify the aggregation of quality statistics for sequencing runs from Illumina NextSeq and HiSeq machines. The single, consistent interface for viewing these data helps to quickly and efficiently identify potential issues with sequencing runs, lanes within the run, or with the original samples. Aggregating information into a single system has the added bonus of allowing extraneous meta-data to be gathered automatically and the overall quality trends in sequencing runs to be viewed. Thus, the data production and run-protocol for each machine can be tracked over time and so guide lab-side decision making and problem solving. In contrast to services such as Illumina’s BaseSpace, AlmostSignificant uses local resources, is open-source, and requires no data to be uploaded. While there are other tools with a similar function to AlmostSignificant (e.g. StatsDB; Ramirez-Gonzalez, 2013), AlmostSignificant has the advantage of being built on the Django web-framework making it light-weight and adaptable. AlmostSignificant works in complement with laboratory management systems such as MISO (http://www.tgac.ac.uk/miso/) or BaseSpace Sequence Hub (https://basespace.illumina.com), complementing their lab-side management with a post-sequencing summary. AlmostSignificant has been in production over the last three years and currently tracks over 80 runs covering over 2500 samples.

## 2 Software Description

### 2.1 Interface

The AlmostSignificant interface is designed to be consistent regardless of the data being viewed but still able to show the information pertinent for the task at hand. The interface is separated into 5 different views; the *Run Overview*, the *Run-details View*, the *Project Overview*, the *Project-details View*, and the *Statistics View*.

The *Run Overview* (Figure 1) shows a table of all the runs, with important metrics for the run shown at the top of the page. Additional information for each run can be accessed but clicking the arrow on the right of each row. This additional information allows at-a-glance checking of the top indexes that could not be assigned to samples (indicative of being given the wrong indexes) and bar-plots to identify any pooling inconsistencies and to ensure each sample produced the required amount of data. The *Run-Detail view* shows a list of all reads for all samples of the run. The interface allows output from FastQC or FastQScreen to be viewed in a single tab. Thus the user can obtain an overview of the quality of the current sample at a single glance, rather than having to open multiple files or scroll down a large HTML page.

**Figure 1.**
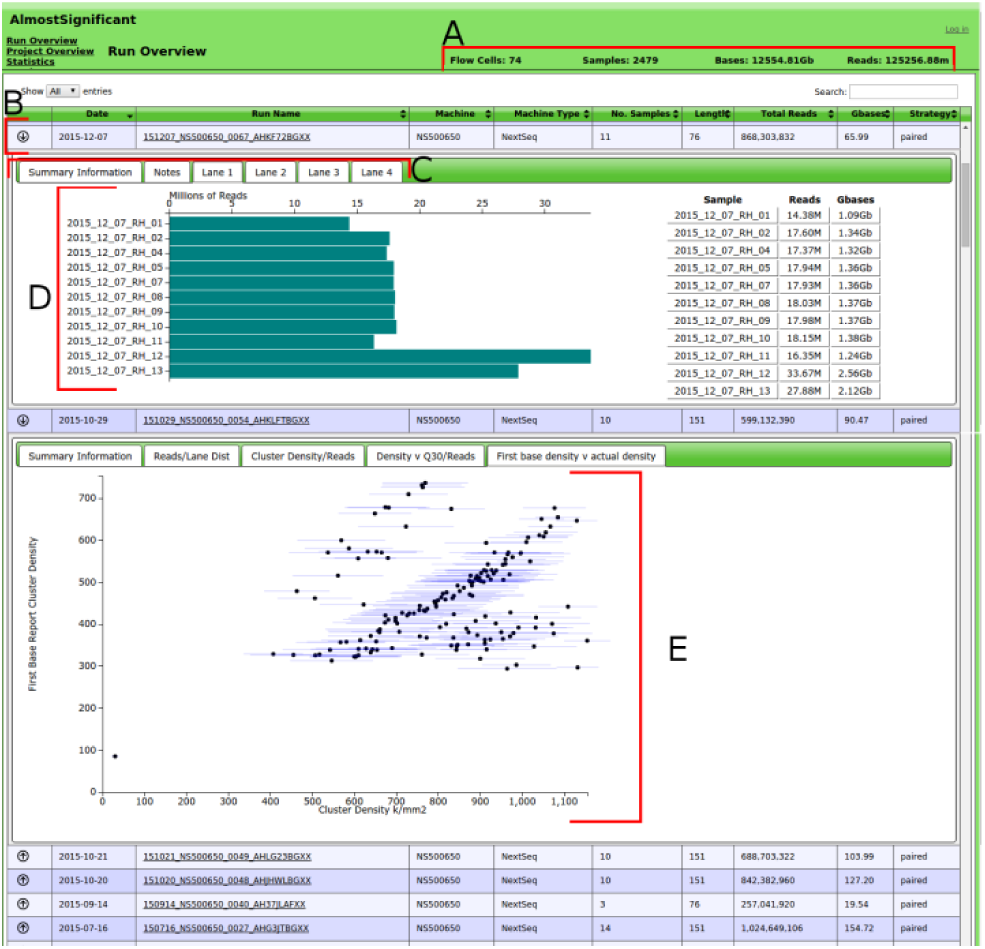
**An example of the interface for** AlmostSignificant showing the overview for all runs, with an expanded row showing the distribution of data produced for all of the samples in a lane. A) A selection of summary statistics relevant to the current view, in this case showing total number of runs and amount of data. B) A row showing information about a single sequencing run. C) Tabs for the expanded drop-down for the selected run, accessed by clicking the arrow on the left of each row. D) Plot showing the data distribution for the samples in the selected lane. E) The relationship between cluster density for a run and the quality of data produced.

The *Statistics View* shows slightly different information to the other views. The rows give a break-down for each length of run for each type of machine used. For example, 100bp HiSeq runs. It shows the number of flow cells that have been run for the run-protocol as well as the mean reads and bases produced for a whole flow-cell or a single lane. Expanding the row shows more details about the run-protocol, including the minimum and maximum output seen in reads and bases, as well as plots comparing the cluster density for the lane to the amount of data produced. Being able to see how the cluster density relates to the data output and quality of the reads allows for optimisation of lab protocol and a target for optimal cluster density.

The *Project View* gathers together information on samples from individual projects, regardless of how many runs they were spread over This allows a simple at-a-glance view of the amount of data produced for each project, similar to the way the *run overview* does for runs. The additional details accessed by clicking the arrow shows the number of times each sample has been run. The project name gives access to a *Project-detail View* showing a summary table similar to the run-details view, but for all samples associated with the project regardless of which run they were sequenced on. This gives an easy way to view the samples for a project without having to visit views for multiple runs.

### 2.2 Data Loading

The data loading script dataLoading.py requires access to three folders as input. The first is the processed-run folder generated by the Illumina bcl2fastq program; this provides information about de-multiplexing and sequences with undetermined indexes as well other meta-data. The second input is the original run folder that is output from the sequencing machine (including sample sheet); this provides information on the run such as the cluster densities and other sequencing-run related meta-data. The third input is a single folder containing all of the quality information output by FastQC and/or FastQScreen. This folder can contain sub-folders but the names for the files should match the fastq files found in the processed run folder. %Optional arguments include selecting whether the run is a HiSeq or NextSeq run (defaults to nextseq) and a flag to tell the script to identify the undetermined indexes found in the Undetermined Index files produced by the sequencing machine.

### 2.3 Requirements

AlmostSignificant requires python 2.7+ and django 1.8+. For optimal usage, the illuminate python package (available via pip) should be installed, and pdflatex should be available from the command line. AlmostSignificant can be run on any standard Linux desktop machine.

## Acknowledgements

We would like to thank Dr Nicholas Schurch (Barton Group) for help with code, and Dr Tom Walsh for computing support.

### Funding

This work was supported by the Wellcome Trust Strategic Award 098439/Z/12/Z.

*Conflict of Interest:* none declared.

